# Ultra-fast and Efficient Network Embedding for Gigascale Biological Datasets

**DOI:** 10.1101/2025.06.18.660497

**Authors:** Jianshu Zhao, Jean Pierre Both, Rob Knight

**Affiliations:** Department of Pediatrics, School of Medicine, University of California, San Diego, California, USA; Independent Researcher, Palaiseau, France; Department of Computer Science & Engineering, University of California, San Diego, California, USA; Halıcıoğlu Data Science Institute, University of California, San Diego, California, USA; Shu Chien-Gene Lay Department of Bioengineering, University of California, San Diego, California, USA; Center for Microbiome Innovation, Jacobs School of Engineering, University of California San Diego, La Jolla, CA, USA

**Keywords:** Graph embedding, Sketching, MinHash, hashing, biological network, link prediction, node classification, Generalized SVD

## Abstract

Graph/network representation learning (or graph/network embedding) is a widely used machine learning technique in industry recommending systems and has recently been applied in computational biology. Popular network representation learning algorithms include random walk and matrix factorization methods, but they do not scale well to large networks. To accommodate the fast growth of real-world network datasets, especially biological datasets, we engineered and improved several network embedding algorithms via intensive computational optimization (e.g., randomized-generalized singular value decomposition/SVD, efficient sketching via ProbMinHash including edge weights) and parallelization to allow ultra-fast and accurate embedding of large-scale networks. We present GraphEmbed, a computer program for scalable, memory-efficient network embedding. GraphEmbed can perform embedding for large-scale networks with several billion nodes in less than 2 hours on a commodity computing cluster. We benchmark it against standard datasets and demonstrate consistent speed and accuracy advantages over state-of-the-art techniques. We also propose centric AUC, a new metric for evaluating link-prediction accuracy in network embedding. It corrects the bias in conventional AUC caused by the highly skewed node degree distributions, which are typically found in real-world networks, especially biological networks. Taken together, GraphEmbed solves a major challenge in large-scale network representation learning for networks in general and biological networks in particular.

## Introduction

Networks or graphs (we use graph and network interchangeably here) are a powerful way to represent data, especially biological data, where data points are connected. A standard approach to computing on networks or graphs is to transform such data into vectorial data—known as graph or network embeddings—to facilitate similarity search, clustering, visualization and community detection^1, 2^. In a network embedding problem, one is given a directed or undirected network and an induced similarity (or distance) function between its nodes; the goal is to find a low dimensional representation of the network nodes in some metric space so that the given similarity (or distance) function is preserved as much as possible^2^. Embedding approaches have several potential advantages. Algorithms using embeddings are frequently faster than their counterparts which operate on the original graphs. The learned embeddings are often applicable for downstream analysis, either by direct interpretation of the embedding space or by applying machine learning techniques that operate on vectors. Traditional unsupervised methods, including matrix factorization and random walk are based on either singular value decomposition (SVD) or SkipGram, which can be computationally inefficient due to memory and run time constraint.

Graph-sampling based techniques design specific models to learn node embeddings from pairs of nodes sampled from an input graph. DeepWalk^3^ and Node2vec^4^ first sample pairs of nodes from an input graph using random walks, and then feed them into a SkipGram-alike model^5^ to output node embeddings. LINE directly samples node pairs from an input graph considering specifically the 1st- and 2nd-order node proximity^6^. VERSE was recently proposed as a generalized graph-sampling based embedding learning framework that preserves a pre-selected node similarity measure^7^. However, because the embedding learning process of the techniques from this category usually relies on Stochastic Gradient Descent (SGD), a large number of node pairs are often sampled and used to ensure the quality of the learned node embeddings (convergence of the SGD algorithms). This requires significant computational resources, CPU in particular.

In addition to graph-sampling-based techniques, factorization-based techniques apply matrix factorization to a high-order proximity/adjacency matrix of an input graph to output node embeddings. GraRep preserves the top k -order node proximity by factorizing the top k-order transition matrices^8^; HOPE investigates several different node proximity measures and uses a generalized SVD to factorize the corresponding high-order node proximity matrices^9^; NetMF was recently proposed as a generalized matrix factorization framework unifying DeepWalk, Node2vec and LINE^10^. Due to expensive matrix factorization operations, the techniques from this category suffer from computational challenges, especially memory. These computational challenges hinder the application of many of those techniques on large-scale networks. In this paper, we focus on unsupervised network embedding, where graphs have no additional node information. We will not discuss graph neural network-based network embedding systems include GCN, GAT and GraphSAGE^11, 12, 13^ because these algorithms usually rely on node attributes/labels, as well as other supervised information.

Real-world networks often have edge attributes, such as positive edge weights, that can reflect the strength of interactions between nodes. Most current network embedding schemes ignore edge weights but ideally network embedding methods should not only be able to capture the (binary) information representing links between nodes, but also relative importance of those links^14^. Taking social networks and protein-protein interaction networks as examples, the edge weights can carry more information on friendship importance and protein-protein interaction strength, respectively. Computational methods for large and sparse networks have received considerable attention in recent years, e.g., random projection-based methods and randomized SVD-based methods have been applied in large-scale network embedding^15, 16^. However, those methods only support binary information between nodes (or simply unweighted edges).

The embeddings learned from the above techniques also face computational challenges when being used in downstream tasks, especially tasks that require similarity computation between embedded node vectors. A typical example is the link prediction problem, in which we try to predict potential missing links between pairs of nodes in a graph to infer possible connections between nodes (e.g., Protein-Protein interactions). Similarity must be computed between all pairs of disconnected nodes in the embedding vector space, then node pairs must be ranked according to their similarity. In the current literature, most of the existing graph embedding techniques measure node proximity using cosine distance (or dot product after normalization). However, cosine distance is known to be slower than Jaccard distance, because dot product with many dimensions is a limiting step^17^. Hashing-based methods such as NetHash and NodeSketch on the other hand, avoids SGD and matrix factorization via sketching algorithms^18, 19^. ProbMinHash,

a MinHash-like sketching algorithm can be used to compute Jaccard-like distance 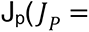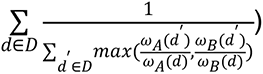 instead of cosine distance between node vectors^20, 21^. MinHash has been widely used for set similarity estimation in large-scale datasets including documents and genomes^22, 23, 24^. NodeSketch showed significant improvement for both CPU time and memory consumption via ProbMinHash. Moreover, it is possible to further optimize the computation of ProbMinHash to be even faster^21^. However, there are no standalone and parallelized implementations for NodeSketch, limiting its application in solving real-world large-scale network embedding problems despite its computationally efficiency.

Benchmarking network embedding algorithms for downstream tasks is another important topic. For example, link prediction or node classification tasks are the main applications in industry. A popular metric to evaluate the link prediction accuracy of network embedding algorithms is AUC (area under the receiver operator characteristic curve, or simply AUC-ROC)^3, 4, 25^. AUC has been widely accepted as a universal benchmark and a downstream task for representation learning. However, its validity is seldom questioned. When link prediction performance is measured according to localized metrics, there is a significant drop in quality^26^. AUC in link prediction is used to benchmark algorithms^27^; used to evaluate new techniques^28^; and draw conclusions about properties of ML algorithms^25^. However, AUC for link prediction by node embedding methods leads to incorrect conclusions about quality^26^. Therefore, a new link prediction accuracy benchmark metric is urgently needed. For node classification accuracy, the F1 score, a widely used metric of precision, is still the gold standard. Using edge weights or other attributes can greatly improve node classification accuracy (F1 score for example), especially for difficult multi-label classification tasks^14^. However, most current embedding algorithms only support unweighted networks due to easier implementation.

Graph embedding has also been applied in computational biology recently, for example, protein-protein interactions prediction^29^, and inferring gene regulatory networks from single-cell transcriptomic data^30^, mass spectrometry data^31^ and other biomedical network datasets^32^. Real- world network and biological network commonly follow an approximately scale-free power-law distribution, meaning that the probability that a node in the network interacts with k other vertices decays as a power-law distribution. Intuitively, scale-free networks contain a few hubs that are connected to many other nodes and many nodes with one or only a few connections. This degree distribution creates a challenge for evaluating the accuracy of network embedding algorithms, for example in the case of link prediction tasks. Recent progress such as *VCMPR@k* ^26^, studied this problem but still, the skewed node degree problem in link prediction tasks remains unsolved (see Methods & Materials section for details).

Here we implemented and engineered an improved network embedding framework that boosts the performance of network embedding algorithms. We also solved the skewed degree distribution problem in link prediction tasks and unweighted edge problem in node classification tasks. We showed that our implementation is on average 60-100 times faster than other popular network embedding algorithms and can scale to networks with up to 3.5 billion nodes. The link prediction metric we proposed, centric AUC, can provide a new standard for link prediction accuracy evaluation for various network embedding algorithms.

## Results

### Centric AUC vs AUC and *VCMPR@k*, Precision and Recall for Link Prediction

We first analyze node degree distribution for some real-world graphs and how it impacts *VCMPR@k*. With an edge deletion probability 𝑝 of 0.2, the binomial distribution of a random graph implies that nodes of degree less than 4 will contribute 0 with probability at least 0.41. Degree quantiles of the Blog, Amazon and Dblp datasets showed we can expect around 10% of nodes to have no edge deleted for the first graph, and around 50% for the second and third (Figure S1A, B and C). Therefore, 12%, 39% and 41% of nodes that do not contribute to *VCMPR@k* for the Blog, Amazon and Dblp networks respectively. We then compute centric AUC, AUC and *VCMPR@k* for those networks via GraphEmbed. The Amazon and Dblp datasets have a similar centric AUC with global AUC in sketching-based embeddings (Table 1). The BlogCatalog dataset has a smaller centric AUC value compared to traditional AUC but in a less pronounced way (e.g., smaller value and standard variation) than that of *VCMPR@k* (Table 1). In both HOPE and sketching embeddings, AUCs are close to 1 while *VCMPR@k* are close to 0, both showing extreme values due to skewed degree distribution (Table 1). We conclude that centric AUC is more versatile for real world graphs with skewed edge distribution and thus should replace AUC as the standard metric for accuracy evaluation of link prediction tasks, a key application of network embedding algorithms.

**Table 1.**
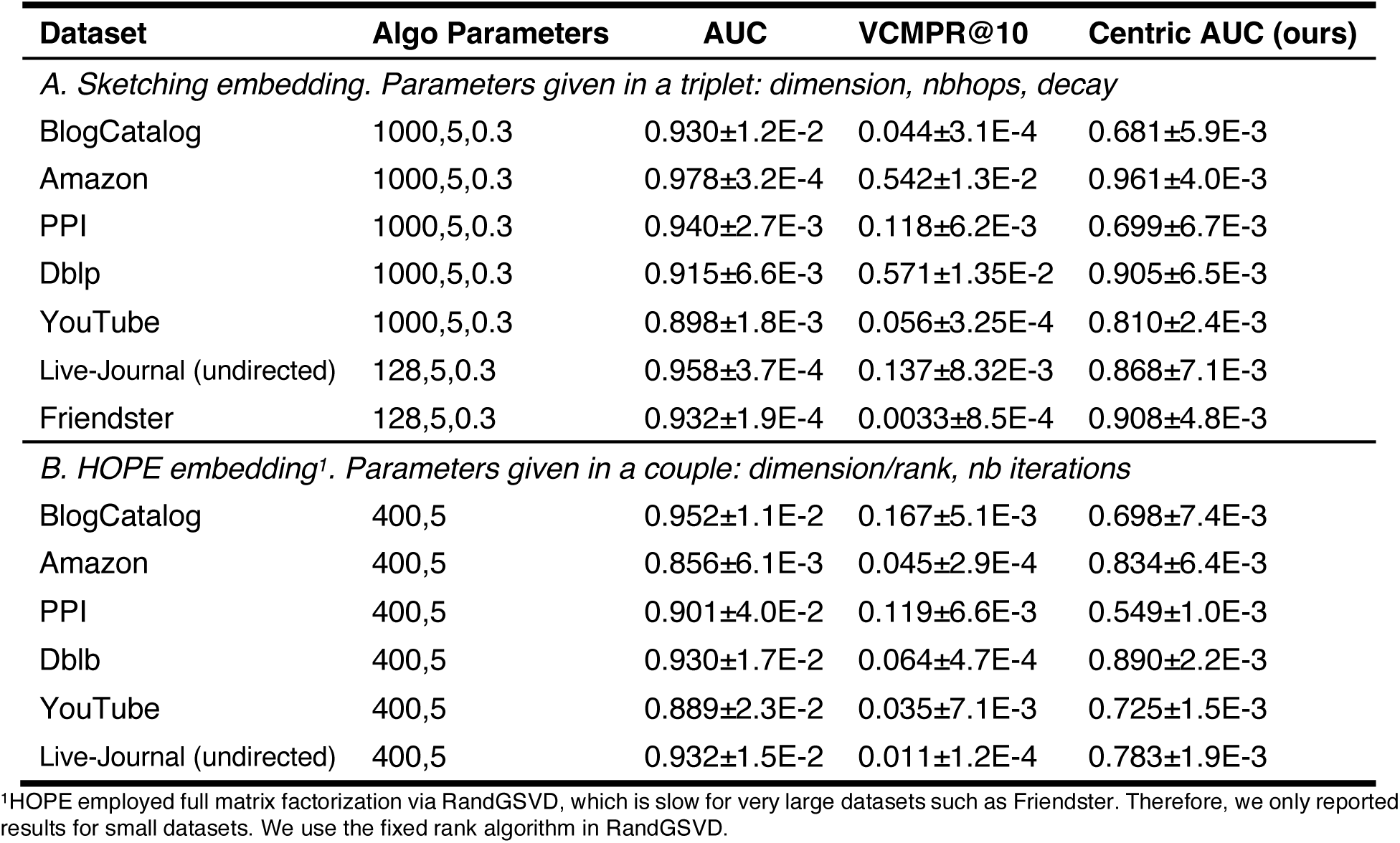
Comparison of AUC, *VCMPR@10* and Centric AUC for link prediction tasks (unweighted network datasets). Dataset information can be found in **Table S1**. Each estimate is the average of 5 independent runs. 2 improved graph embedding algorithms implemented in GraphEmbed, Sketching and HOPE were evaluated.

We then studied the effects of hyperparameters on those link prediction metrics. For HOPE precision mode, hop (i.e., iteration around a node) and precision (RandGSVD precision) had almost no effect on AUC but small effect on *VCMPR@10* and centric AUC (Figure 3C, D and E). For the sketching algorithm, we observe a clear convex surface for each metric (Figure 4C, D and E) with respect to two hyperparameters, hop and decay weight ⍺. For precision and recall in sketching mode, a similar convex surface was observed, suggesting optimal hop and ⍺ for link prediction task (⍺=0.3 and k=5) (Figure S3A and B). Those results suggest that node embeddings are often task-specific and should be appropriately generated by tuning parameters for individual network analysis tasks, see also discussion in Yang et.al. (2022)^33^. For HOPE with RandGSVD in precision mode, the effect of RandGSVD precision and hop on link prediction recall and precision is small (Figure S4A and B). However, we observed that *VCMPR@k* and centric AUC are more sensitive than AUC (Figure 3C, D and E). In the rank mode, the effects of hop on all 3 metrics are small but still *VCMPR@k* and centric AUC are more sensitive than AUC (Figure S5).

**Figure 1.**
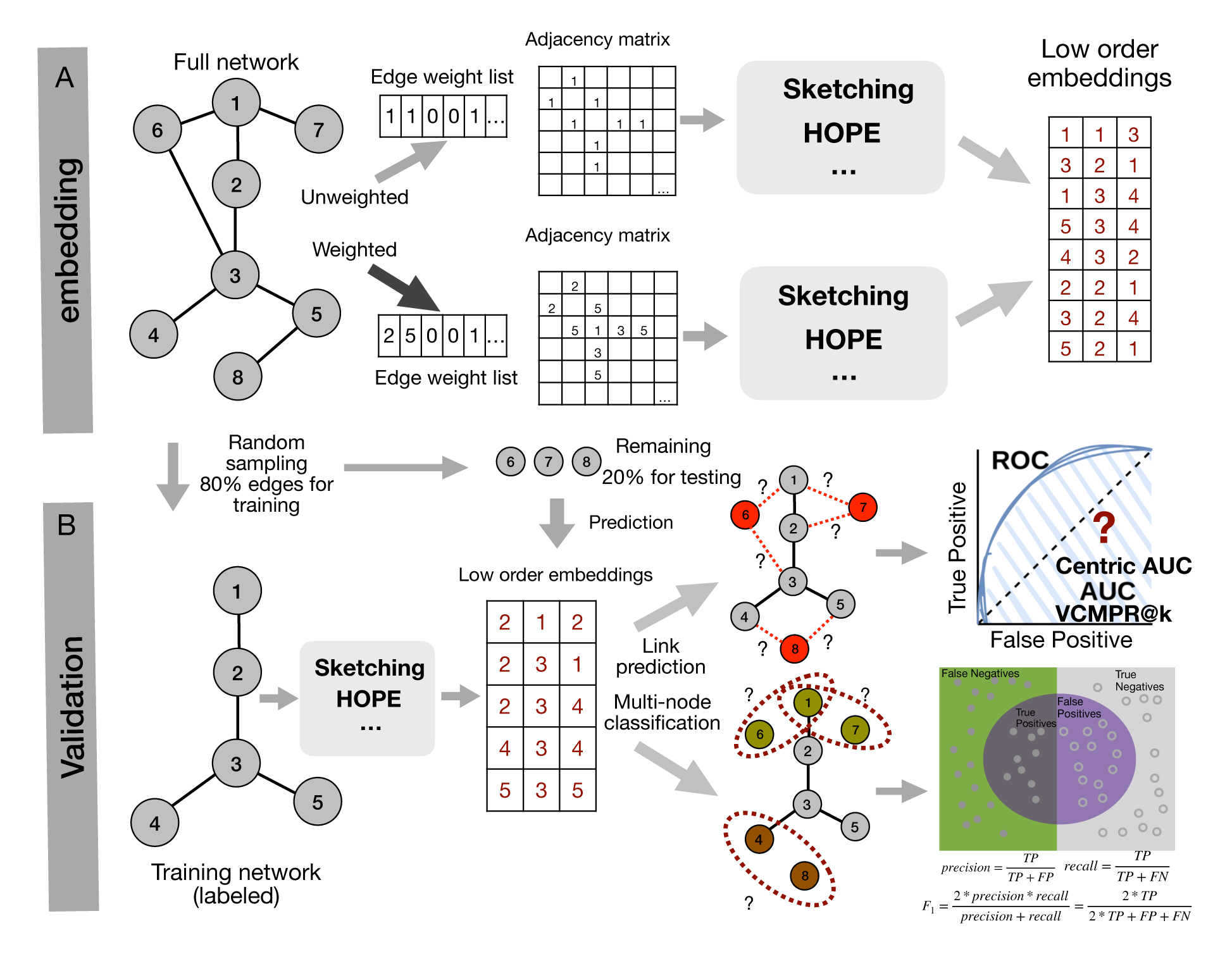
Schematic overview of GraphEmbed. Upper panel is the embedding mode, in which only the actual embedding process will be performed. HOPE algorithm or sketching algorithm can be used by default. Bottom panel is the validation mode, in which detailed accuracy benchmarks for link prediction and node classification can be performed. The subsampling ratio for training and testing can be easily tuned.

**Figure 2.**
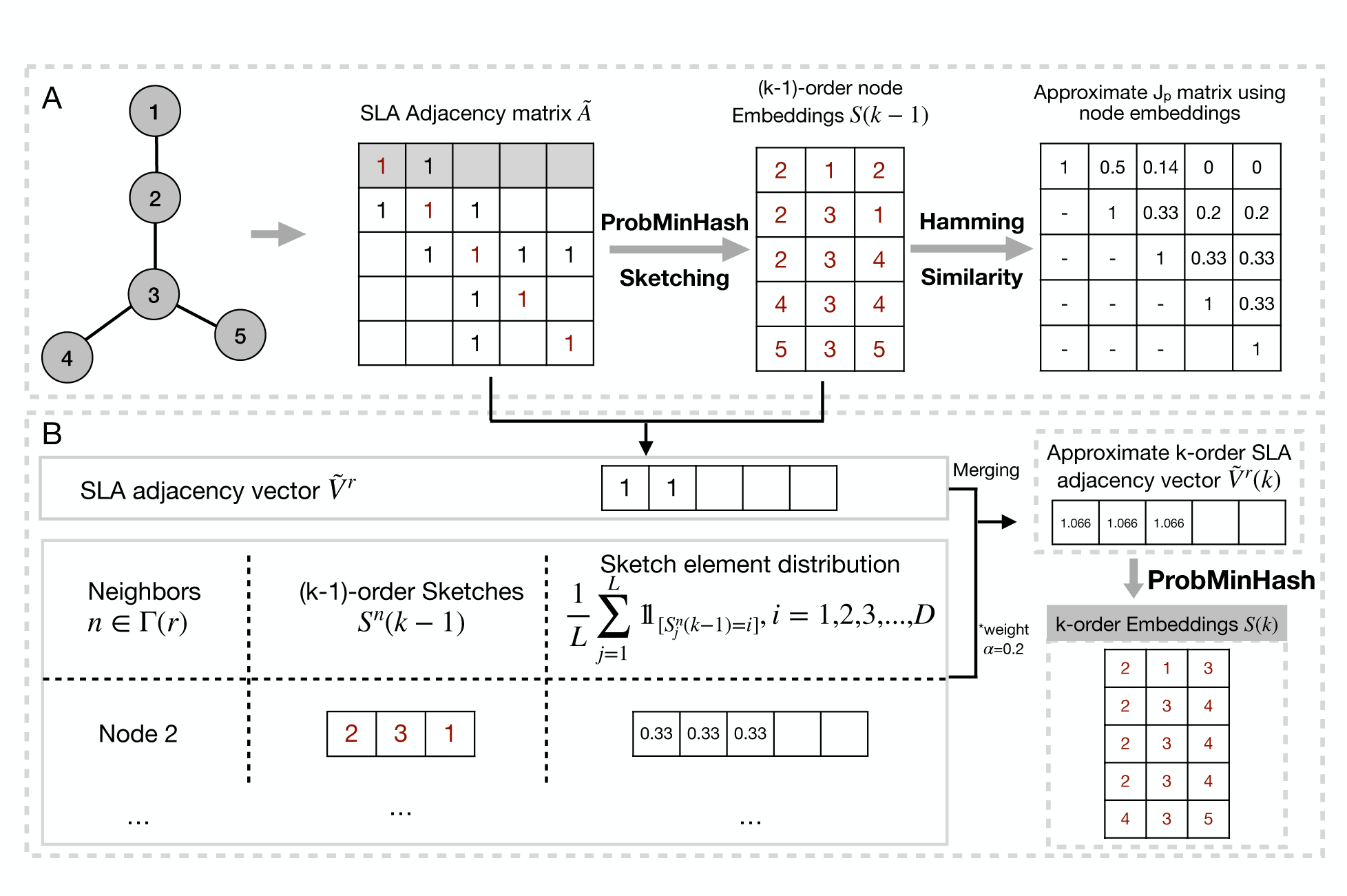
Overview of GraphEmbed sketching algorithm, an improvement over NodeSketch. (A) Low-order node embeddings can be obtained by sketching the Self-Loop-Augmented (SLA, that is the diagonal in red) adjacency vector of each node, where the sketch length is set to 3. The SLA adjacency vector and the corresponding embeddings of node 1 were highlighted as an example. Then J_p_ can be efficiently approximated by computing the hamming similarity between node embeddings, which equals the collision probability of hashes in the sketch. (B) High-order node embeddings (in this case k=3) via recursive sketching. The detailed recursive sketching process for node 1 based on the SLA adjacency matrix and k-order node embeddings was used as an example. The approximated k-order SLA adjacency vector 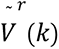 by summing the SLA adjacency vector of node 1 and the sketch element distribution in (k-1)-order embeddings of all nodes 1’s neighbors (node 1 has only one neighbor node 2 in this case) in a weighted manner (weight is alpha=0.2). Then, the k-order node embeddings were obtained by sketching the approximated k-order SLA adjacency vector 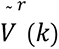 via ProbMinHash.

**Figure 3.**
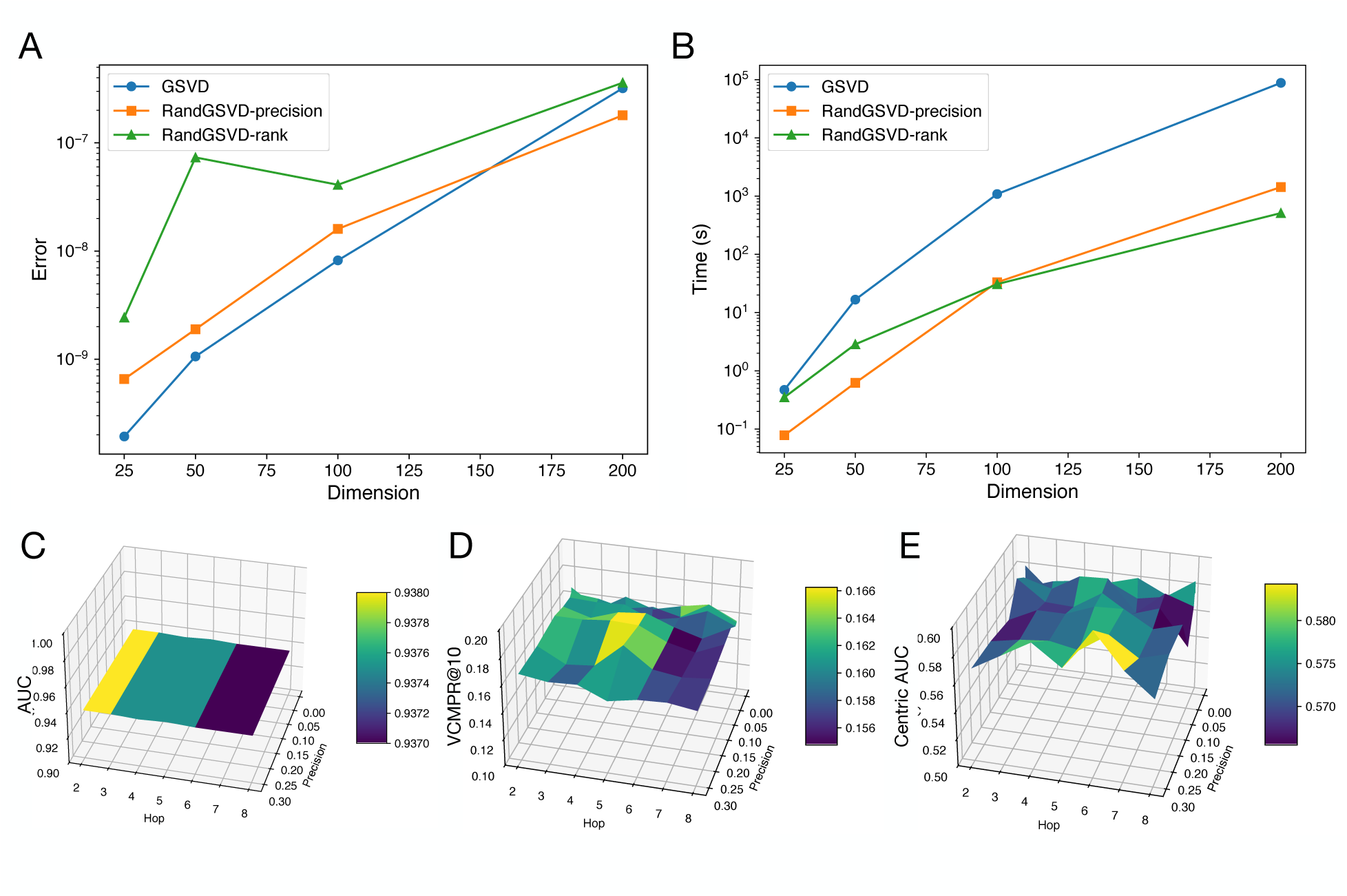
Accuracy (A) and speed (B) benchmark for Randomized Generalized SVD against original generalized SVD for simulated data with increasing dimension. Effects of HOPE RandGSVD precision and hop on link prediction accuracy in the fixed precision algorithm for traditional AUC (C), *VCMPR@10* (D) and centric AUC (E), respectively. Maximum rank was set to 128. The BlogCatalog dataset was used for the plot.

**Figure 4.**
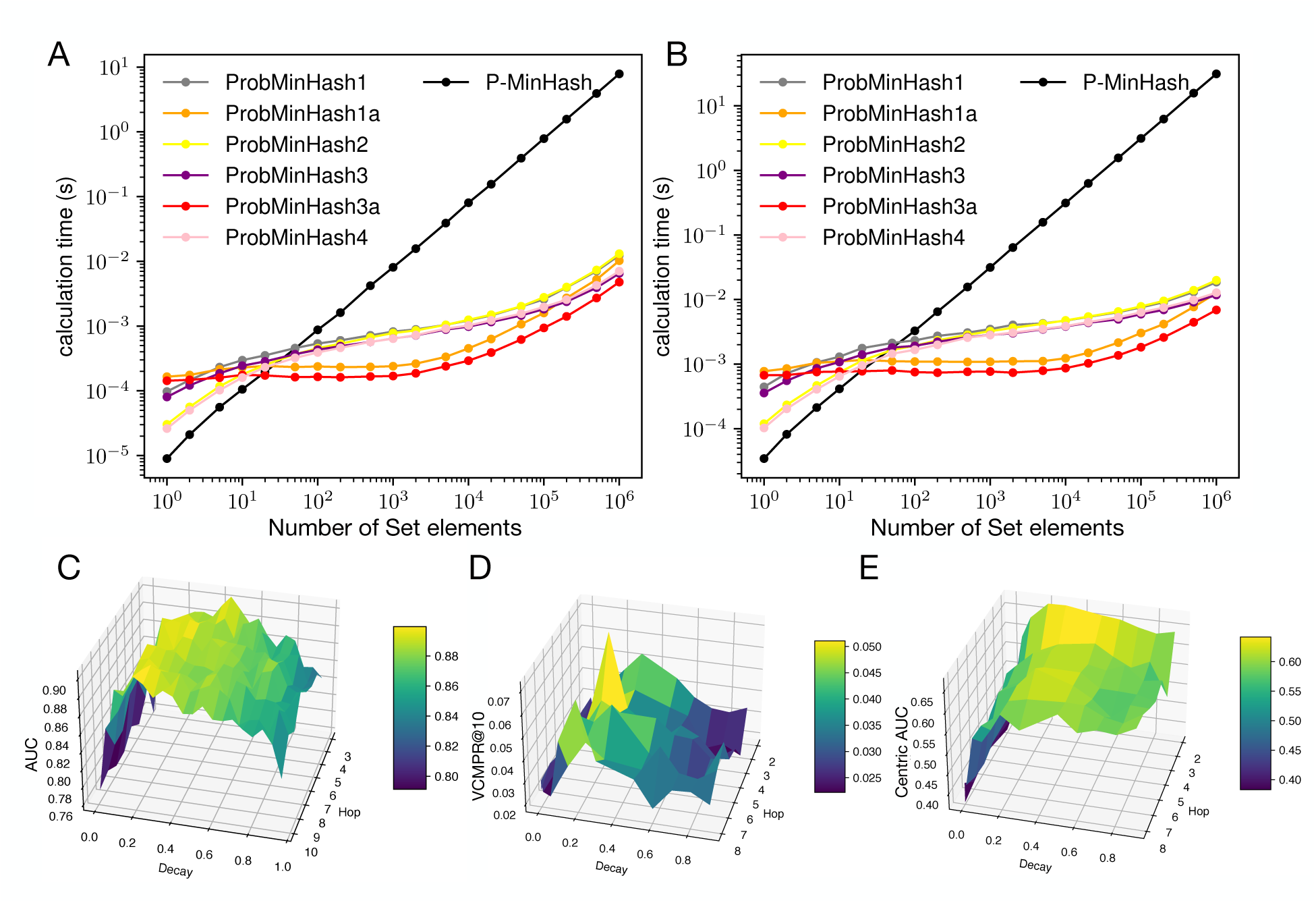
ProbMinHash algorithm benchmark with respect to number of set elements (or dimension in adjacent vector) and sketch size m=1024 (A) and m=4096 (B). The weight of set elements was simulated via an exponential function 𝑤(𝑑) ∼ 𝐸𝑥𝑝(1). The effect of decay (𝛼) (x- axis) and number of iterations or hops (y-axis) in GraphEmbed sketching on AUC (C), *VCMPR@10* (D) and centric AUC (E) (z-axis). The underlying dataset is BlogCatalog for (A), (B) and (C).

### Link Prediction Precision and Recall

Benchmarks of our HOPE implementation and ProbMinHash-based sketching showed that GraphEmbed is in line with the original implementation for various datasets (Table S2). Based on precision and recall, GraphEmbed (sketching) beat other sketching-based or lean-to-hash based embedding methods consistently across all benchmark datasets and can be more accurate than some of the matrix factorization or random walk-based methods for the Dblp and PPI datasets (Table S2) while being much faster (see next section). However, GraphEmbed (HOPE) is slightly less accurate than other matrix factorization methods such as VERSE and node2vec in terms of precision and recall, the later methods are overall more than 30 times slower than our HOPE implementation and cannot scale even for medium size networks (Table 2). We did not use AUC for benchmarking link prediction accuracy in this case because all datasets mentioned above have skewed node degree distribution.

**Table 2.** Embedding speed benchmark (in seconds) and the average speedup over state-of-the-art matrix-factorization and random-walk based methods. Note that GraphEmbed (sketching) multi-threaded implementation can be more than ∼60 times faster than GraphEmbed (HOPE) implementation. Speedup within the category was calculated using GraphEmbed method (HOPE or sketching) as the reference point, respectively. Speedup across categories was calculated using GraphEmbed (sketching) as the reference point. – indicates that the algorithm cannot produce results in less than 3 days.

| Methods | Blog | PPI | Dblp | Wiki | YouTube | Speedup<br>(within<br>category) | Speedup<br>(across<br>categories) |
| --- | --- | --- | --- | --- | --- | --- | --- |
| DeepWalk | 3143.2 | 1145.0 | 4421.3 | 1134.1 | 698509.1 | 117.9x | 7840.3x |
| LINE | 1994.7 | 2001.6 | 2322.1 | 1776.0 | 24736.8 | 58.4x | 3383.6x |
| node2vec | 1012.3 | 335.5 | 472.0 | 1208.45 | - | 36.1x | 2400.6x |
| VERSE | 983.1 | 181.9 | 1025.4 | 259.3 | 215432.0 | 31.8x | 2114.7x |
| NetMF | 453.0 | 111.2 | 209.7 | 631.9 | - | 17.8x | 183.7x |
| GraRep | 3019.5 | 321.1 | 8912.3 | 408.2 | - | 90.6x | 6024.9x |
| HOPE | 251.4 | 97.7 | 325.0 | 83.6 | 15265.3 | 7.0x | 332.5x |
| **GraphEmbed-HOPE<sup>1</sup>** | **42.1** | **13.0** | **54.8** | **11.1** | **2483.4** | **1.0X** | 66.5x |
| NetHash | 701.3 | 183.9 | 29.1 | 131.6 | 11432.9 | 341.2x | 341.2x |
| KatzSketch | 189.6 | 19.1 | 232.1 | 35.4 | 8949.2 | 243.4x | 243.4x |
| NodeSketch<sup>1</sup> | 56.9 | 7.4 | 9.2 | 15.2 | 2513.7 | 59.6x | 59.6x |
| **GraphEmbed-Sketch (1)** | 13.5 | 4.4 | 4.3 | 5.4 | 843.9 | 28.2x | 28.2x |
| **GraphEmbed-Sketch (64)<sup>2</sup>** | **0.2** | **0.3** | **0.1** | **0.4** | **28.4** | **1.0X** | **1.0X** |

<sup>1</sup>Single-threaded only; <sup>2</sup>Parallelized for all steps. 64 threads were used.

### Node Classification Accuracy

GraphEmbed sketching algorithm outperforms all sketching baselines except for the Wiki dataset, where baseline KatzSketch is slightly better than GraphEmbed (Table S3). However, KatzSketch involves expensive matrix multiplication/inversion operations to compute the Katz index while GraphEmbed simply relies on ProbMinHash and showed a 243x speedup over KatzSketch on average (Table 2). Among classical network embedding methods, NetMF achieves the best performance on Blog, Wiki and DBLP, while DeepWalk and VERSE are the best ones on POI and YouTube datasets, respectively (Table S3). GraphEmbed sketching shows comparable performance to these best-performing baselines; it has better results on the PPI, Wiki and DBLP datasets, and slightly worse results on Blog and YouTube datasets (Table S3). However, GraphEmbed sketch is far more efficient than these baselines, i.e., 183x, 7840x and 2114x faster than NetMF, DeepWalk and VERSE, respectively (Table 2). The HOPE implementation in GraphEmbed showed similar accuracy with the original implementation in terms of both Micro-F1 and Macro-F1 (Table S3), and 7X faster than original HOPE (Table 2). For weighted network, node classification accuracy is significantly improved by considering weight (Table S6) despite that link prediction accuracy is compromised (Table S5).

### Embedding Gigascale Networks in Approximately One and A Half Hours

To compare our ProbMinHash implementation with reference implementations, we simulated weighted vectors and tested the sketching speed. We found that our implementation is about ∼1000 times faster than the original ProbMinHash in NodeSketch for vectors with a million elements or more, and performance is comparable for fewer elements (Figure 3a and b). For sparse networks (many entries are empty in adjacency vectors), we did not expect such a huge speedup because the second pass of data stopped earlier for both original ProbMinHash and our implementation (Figure S2). Next, we compared GraphEmbed HOPE with other matrix factorization or random walk-based methods. First, we compare the accuracy and running time of the limiting step in HOPE, randomized-generalized SVD, with that of traditional SVD. We showed that RandGSVD achieves accuracies almost as good as those of the traditional GSVD but up to dozens of times faster, especially when the dimension is large (Figure 3a and b). Overall, our HOPE implementation in GraphEmbed is about 7 times faster than the original HOPE, which utilizes Jacobi-Davidson partial SVD, due to computational optimization provided through RandGSVD (Table 2, single thread). We observed a ∼20 times speedup with our HOPE implementation if parallelized with 64 threads. For small and dense network datasets GraphEmbed sketching method is on average ∼60 times faster or more than other hashing-based methods (Table 2) and scales well with respect to the number of computing threads (Figure S6). Detailed CPU and memory usage analysis for test datasets showed that the sketching algorithm is highly efficient (Figure S7). For large-scale sparse networks (unweighted), GraphEmbed sketching took ∼1.5h for Hylerlink2012 dataset and is as fast as cutting-edge network embedding algorithms such as SketchNE, that aiming at sparse networks only and is 5 to 10 times faster than other sparse network embedding algorithms (Table S4). However, the memory consumption for processing 3.2 billion nodes in parallel is approximating 1.8T, slightly larger than sparse-signed randomized SVD in SketchNE (1.5T) for the same dataset due to ProbMinHash sketching algorithm (Table S4).

## Discussion

In this paper, we provided a fast and efficient network embedding system that is standalone and can be easily deployed to commercial platforms for large-scale (biological) network embedding. We also provide embedding quality evaluation (e.g., link prediction accuracy and node classification accuracy) so that we can directly compare the performance of various network embedding algorithms. In the case for link prediction tasks, the new metric, centric AUC we proposed showed interesting properties compared to the traditional AUC and *VCMPR@k* for real- world networks. We suggest that centric AUC should be used instead of traditional AUC because it considers highly uneven node degree distributions, as typically found in real-world networks. We showed that in many of our benchmarks for real-world networks, AUC is very close to 1 while *VCMPR@k* is very close to 0 and centric AUC is in the middle. Centric AUC is versatile and has a less extreme distribution on different datasets (Table 1). It has been pointed out that the global “dense metric” of AUC is not attuned to the sparse ground truth, which was the motivation for *VCMPR@k*26. However, the *VCMPR@k*, for small k, is an averaged “local” score, and is more appropriate for sparse ground truth without considering from the global “dense metric” perspective. It was suggested that *VCMPR@k* can only be used in networks that are “globally sparse, yet locally dense” (often measured as low global density and high clustering coefficients)^26^. For example, *VCMPR@k* is not meaningful for the PPI (protein-protein interaction) dataset, which has low clustering coefficients, as do many other biological networks. Our centric AUC method can be considered as a mixture of AUC and *VCMPR@k* and can be applied to any type of networks because the edge sampling is proportional to the degree of nodes. It has been pointed out that AUC is deeply flawed because it creates a data-dependent reweighting of false positives vs. false negatives^34^. Therefore, precision and recall measures in a global manner should be used^35^. We also implemented global precision and recall according to^36^ but it is extremely computationally expensive because it requires computing the whole (number of nodes) matrix similarity between embedded nodes and sorting them to compare to the deleted edges.

This method becomes impractical at a few thousand nodes or above (e.g., the YouTube dataset) even though recall has been proven to be an unbiased metric towards different node degrees^37^. Centric AUC employs a degree-specific sampling strategy and is thus more scalable for large networks (Table S7). Another implicit bias related to node degree was also found recently and the author proposed a degree-corrected link prediction task that more effectively trains graph machine-learning models by reducing overfitting to node degrees^38^. This degree-corrected idea is different from our centric AUC and it also yields smaller values than traditional AUC for various benchmark datasets. A node degree specific sampling strategy was used in our implementation instead of sampling negative edges with the same degree bias as positive edges^38^. However, it was shown in the degree-corrected sampling paper that for real-world networks with high degree heterogeneity, naive degree-based methods (our method) can achieve near-optimal performance^38^. Since there is no need to sample both negative edges and positive edges, naive degree-based methods can be significantly faster and scale to large networks. The degree-corrected *VCMPR@k* also yields much larger value than the original *VCMPR@k* and smaller values than AUC, consistent with centric AUC.

Our implementation of HOPE adopted Adamic Adar representation/index of network, which is the most widely used set similarity metric in social networks. The index was proposed as an alternative to Jaccard index for the problem of predicting links, such as friendship or co- authorship, in social networks^39^. Recent work on this index showed that we can approximate it via a new sketching algorithm called DotHash^40^, in which one can estimate the set intersection cardinality (or shared neighbors between two nodes) without bias by computing the dot product of between two DotHash sketches. Then the Adamic Adar index can then be efficiently approximated. The current limiting step in our HOPE implementation for large-scale networks is the singular value decomposition (SVD) despite our efforts to speed it up via RandGSVD. Single-pass randomized SVD (key step of RandGSVD) can be further improved as in Yu et.al. 2017^41^, so that a new iterative QB approximation and the power iteration scheme can be incorporated into the single-pass SVD ^42^ via window-based power iteration^41, 43^, which will further improve accuracy but will require 2 passes. A dynamically shifted power iteration technique can also be applied to improve the accuracy of the randomized SVD. Sparse-signed randomized SVD can also be employed in HOPE for better scalability and less memory constraint, e.g., via the sparse Personalized PageRank proximity measurement between nodes ^44^.

The sketching-based network embedding scheme (ProbMinHash) in our system is as fast as the cutting-edge network embedding algorithm such as SketchNE ^45^ or STRAP ^44^, which employed sparse-sign randomized streaming SVD^42, 46^. However, sparse-sign randomized SVD works well only if eigenvalues are well separated and that is only expected for sparse networks^47^, not dense networks, which also helps explain why SketchNE is slower than other sketching-based embedding methods and less accurate for small and dense dataset such as BlogCatalog^45^. Therefore, SketchNE cannot be used for regular networks. Additionally, SketchNE only considers unweighted network data. However, SktehNE is memory efficient than other large-scale network embedding algorithms, including GraphEmbed because the Graph Based Benchmark Suite (GBBS) graph representation format^48^ is a better graph data compression format than compressed row representation (CSR) format^49^, which GraphEmbed relies on. We plan to use GBBS graph representation format in future to reduce the memory requirement for graph processing.

Our ProbMinHash-based sketching algorithm for computing Jaccard-like distance (J_p_) is highly efficient and thus can scale to large networks, either sparse or dense, with weighted edges or unweighted edges. Similar approaches such as SimHash^50^, which is also a locality sensitive hashing and approximates cosine similarity, have also been applied in large-scale attributed network embedding^51^. Another interesting perspective regarding the actual similarity metric for measuring distance between node representation is that J_w_ can be calculated via BagMinHash^52^ or a faster alternative DartMinHash^53^, instead of ProbMinHash or D^2^HistoSketch (which approximate J_p_), because the weight in recursive sketching can be considered. Both J_p_ and J_w_ are metric distances and the MinHash algorithm approximating them are all locality sensitive hashing. It would be interesting to also implement DartMinHash in GraphEmbed in the future. However, MinHash algorithms for simple Jaccard, which are generally much faster^23, 54, 55, 56^, cannot be used under the recursive sketching scheme because element weight is not considered in those algorithms (there will always be a weight for the approximate k-order adjacency vector, Figure 2).

GraphEmbed does not provide implementation or benchmark for random-walk based network embedding because it has been shown that SkipGram with random-walk is theoretically connected to matrix factorization-based methods: SkipGram with random-walk is essentially performing implicit matrix factorization^10^. Those methods were compared with NetMF in previous work, which we frequently benchmarked GraphEmbed against. Additionally, GRAPE provided a convenient network processing and random walk embedding system and is also in Rust^57^. The current GraphEmbed system also does not consider deep structural information of networks, which can be important since real-world networks contain global structures. Structure Deep Network Embedding (SDNE) first considers the high nonlinearity in network embedding and proposes a deep autoencoder to preserve the first- and the second-order proximities jointly to preserve the deep network structure^58^. However, SDNE is very difficult to scale to large networks since the limiting step is still stochastic gradient descent. A specific case for structural information of networks is the bipartite networks, in which 2 classes of nodes (they are not treated equally) will be jointly modeled so that the bipartite structure can be considered^59^. Bipartite network embedding is another interesting problem that has been carefully analyzed at large-scale^60^ and can be applied for large-scale microbiome data analysis. Recently, it was found that node-specific metrics such as the node local intrinsic dimension (NC-LID) can reflect the local degree of the discriminatory power of an arbitrary graph-based distance function (all above network embedding algorithms use the same graph-based distance function for all nodes)^61^. NC-LID can identify nodes with high link reconstruction errors in Node2Vec embeddings. NC-LID can be used as a choice for designing elastic graph embedding algorithms in which hyper-parameters are personalized per node and/or link and adjusted from some base values according to structural graph measures to effectively improve the quality of network embedding algorithms^61^. Additionally, node LID-aware methods can also help design new sampling strategies for link prediction accuracy evaluation, e.g., node LID-aware centric AUC or *VCMPR@k* . However, designing such an embedding algorithm considering different nodes with different NC-LID values can be challenging because we need to compute NC-LID for each node, which can be computationally expensive and difficult to scale to large networks.

In summary, the GraphEmbed network embedding system GraphEmbed we developed will aid large-scale network analysis and facilitate application in industry-level recommending systems, as well as designing faster and more efficient network embedding algorithms (via the new link prediction metric). The flexible interface and parallelized implementation can be easily tuned to integrate new network embedding algorithms. GraphEmbed will save a vast number of computational resources (in terms of CPU hours) and thus facilitate knowledge capture in network biology and more generally, network and machine learning sciences.

## Methods & Materials

The overall schematic overview of the GraphEmbed system can be found in Figure 1. Specifically, there are two modes in GraphEmbed, embedding mode and validation mode. Embedding mode only performs embedding while validation mode also performs embedding accuracy evaluation, including link prediction accuracy evaluation with AUC, centric AUC and *VCMPR@k* and multi-node classification accuracy with F1 score (Macro or Micro, see below). For both modes, 2 embedding options, HOPE and Sketching can be chosen. We applied cutting-edge computational optimization to speed up HOPE and sketching algorithms. We also made GraphEmbed a flexible system so that new embedding algorithms can be easily integrated into this framework without worrying about other implementations (e.g., network representation and processing, embedding performance evaluation). For validation mode, we also provided an option to vary the subsampling ratio of the full network and an option for running a given number of repeated experiments for training and evaluation.

### Precision, AUC, *VCMPR@k* and Centric AUC

First, we revisit the link prediction metric *VCMPR@k* ^26^, which is an alternative to frequently-used AUC and considered an uneven distribution of edge degree. It is defined as the mean over nodes i of 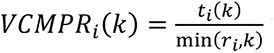 with 𝑟*_i_* being the number of edges deleted from the full graph to the train graph and 𝑡*_i_* being the number of ground truth edges in this list. However, for many real-world graphs, node degrees follow a power law distribution^62^; a large fraction of nodes may have degree much lower than the mean degree. These impacts *VCMPR@k* as 𝑉𝐶𝑀𝑃𝑅*_i_*(𝑘) is not defined for nodes with no edge to delete. However, it is possible to sample the nodes according to node degree distribution for edge deletion. We define centric AUC in the following way: Let 𝑖 be a node, 𝑑*_i_* its degree, 𝑟*_i_* the number of deleted edges incident to 𝑖. After edge deletion, a node 𝑖 has degree 𝑑*_i_* − 𝑟*_i_* and we have 𝑛 − 1 − (𝑑*_i_* − 𝑟*_i_*) potential edges to test. Given a node 𝑖, we first sort the n-1 edges by decreasing prediction of existence of a true edge between nodes 𝑖 and 𝑗. We then define 𝑐*_j_* as the number of true edges seen up to node 𝑗. When exploring the 𝑗-th node, there are 2 possibilities: (1). 𝑗 corresponds to a training edge, if so, we increment 𝑐*_j_* by 1; 2. 𝑗 corresponds to a deleted edge. As our array is sorted, the probability that this edge has a higher ranking than a random edge is simply the ratio of potential edges after 𝑗 to the total number of potential edges 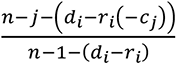. Averaging over deleted edge indexes 𝑗, this defines a centric AUC at node 𝑖, then averaging over uniformly sampled nodes, we have a centric AUC for all sampled nodes. The centric AUC estimation was then compared with original AUC and *VCMPR@k*. Note that centric AUC and *VCMPR@k* works for both directed and undirected networks but the sampling for each node should be direction aware for directed networks.

We also implemented precision and recall according to Lü and Zhou^36^. Specifically, precision and recall were calculated the following equation: 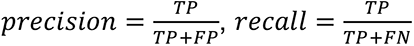, where TP is true positives, FP false positive and FN is false negative. Precision is the proportion of correctly predicted positive links (edges) out of all the predicted positive links while recall is the proportion of correctly predicted positive links (edges) out of all the actual positive links in the network. Since precision and recall works for both directed and undirected networks, the edge deletion is performed as follows: For a directed network edge 𝑖 to 𝑗 and 𝑗 to 𝑖 are deleted/kept together whereas for an undirected network they are treated independently.

### Computational optimization for HOPE: Randomized and Generalized Singular Value Decomposition (RandGSVD) with parallelization

In several network node representation formats in original HOPE (**H**igh-**O**rder **P**roximity preserved **E**mbedding), we implemented 3 of them, Katz index, Personalized PageRank and Adamic Adar^9^. Adamic Adar is the default option since it is widely used in social network link prediction^39^. Instead of the generalized SVD (Jacobi–Davidson type partial SVD) used in original paper^63^, we instead use randomized and generalized SVD (RandGSVD)^64^ for matrix factorizations step in our HOPE implementation, as it works for all matrices, dense or sparse with deterministic error bound under both the spectral norm and Frobenius norm^42, 65^. Specifically, the node proximity measurements problem (multiplication of two polynomials of matrices) under any of the 3 indexes mentioned above can be transformed into a generalized SVD problem^9^. Randomized-generalized SVD is more efficient in time and space than traditional generalized SVD due to randomized SVD algorithm^42^. Furthermore, randomized SVD has also been used for dimensionality reduction tasks (e.g. PCA) in large-scale biological data analysis in a streaming fashion (e.g., single cell RNA sequencing data, microbiome data)^41, 42, 43^ although it exhibits some accuracy loss for high rank matrices^66^. We implemented both versions of RandGSVD, fixed precisions and fixed rank, corresponding to algorithm 2.2 and 2.4 as in Wei et.al. (2021)^64^. The underlying single-pass randomized SVD was implemented according to Halko et.al. (2010)^42^, including both fixed precisions and fixed rank mode. Basic matrix operations were supported via BLAS (Basic Linear Algebra Subprograms) interface. The original HOPE algorithm was designed for directed networks to preserve transitivity^9^ but in our implementation, we provided additional support for undirected networks so that both *VCMPR@k* and centric AUC can be applied to evaluate link prediction performance, which also preserve transitivity for undirected networks.

### ProbMinHash sketching considering edge weights

Our network embedding algorithm based on sketching extends and improves the original sketching algorithm in NodeSketch^18^. Briefly, GraphEmbed sketching learns high-order node embeddings in a recursive manner via ProbMinHash (Figure 2A, B and Supplementary Figure S2). To output the k-order embedding of a node, it sketches approximate k-order Self-Loop-Augmented (SLA) adjacency vector of the node via ProbMinHash, which is generated by merging the node’s SLA adjacency vector with (k-1)-order embeddings of all the neighbors of the node in a weighted manner. One key property of the sketching technique is the uniformity of the generated samples, which serves the foundation of the recursive sketching process. Based on the uniformity property, the sketching process works in the following way. First, for each node r, we compute an approximate k-order SLA adjacency vector 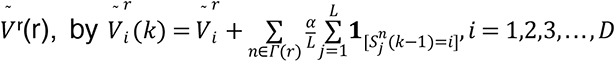 where 𝛤(𝑟) is the set of neighbors of node r, 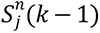 is the (k-1)-order sketch vector of node n, and 𝟏_[cond]_ is an indicator function, which equals 1 when *cond* is true and 0 otherwise (Figure 2B). More precisely, the sketch element distribution for one neighbor n, (e.g., 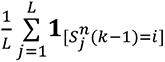) approximates the (k-1)-order SLA adjacency vector of the neighbor, which preserves the (k-1)-order proximity. Subsequently, by merging the sketch element distribution for all the node’s neighbors with node’s SLA adjacency vector, we indeed expand the order of proximity by one and therefore obtain an approximate k-order SLA adjacency vector of the node (Figure 2B). During the summing step, we assign an exponential decay weight 𝛼 to the sketch element distribution, in order to give less importance to high-order node proximity. This is a widely used weighting scheme in measuring high-order node proximity^9^. Finally, we generate the k-order node embeddings 𝑆(𝑘) by sketching the approximate k-order SLA adjacency vector 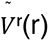 using hash function in ProbMinHash. Note that for the weighted edge network, the SLA adjacency vector of each node will have weight for each of its edges but 1 for itself (diagonal) (Figure 2A), other steps are the same with unweighted edge. We also extend the original NodeSketch algorithm (that only supports undirected network) to allow directed network embedding. All the steps are the same except that the SLA adjacency matrix is not symmetric.

We implemented and optimized the ProbMinHash algorithm as introduced in Ertl (2020) ^21^ and Li et.al. (2022)^67^ as an alternative to D^2^HistoSketch in NodeSketch for the sketch step mentioned above^18, 68^. Specifically, a class of optimized ProbMinHash algorithms were implemented, including ProbMinHash1, ProbMinHash1a, ProbMinHash2, ProbMinHash2a, ProbMinHash3, ProbMinHash3a, ProbMinHash4 and ProbMinHash4a. They all share the same estimation accuracy with original ProbMinHash. Note that both ProbMinHash and D^2^HistoSketch approximate 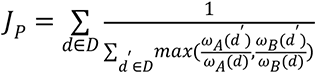, instead of 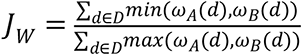 (or min-max similarity)^21^. J_p_ is a generalized min-max similarity index on probability distributions, and it is pareto optimal^67^. See supplementary methods for the comparisons between J_p_ and J_w_. Similar to SuperMinHash, multiple hash functions were used in ProbMinHash so that when there are empty buckets, up to *s* rounds of re-hashing the input with a new hash function *h_i_*. This is sufficiently fast in expectation, since it’s exceedingly rare to have empty buckets. If there are still empty buckets, we fall back to s hashes *h*’ , the *i*th of which maps *all* values into bucket *i* so that it is guaranteed each bucket will eventually be non-empty. Over the first *s* rounds, every element is mapped exactly once to each bucket by explicitly constructing a permutation to control the bucket each element is mapped to in each round^69^. ProbMinHash also tracks for each bucket the best value seen so far, and ensures that for each element, the hash-values for each bucket are generated in increasing order so that the insertion of an element can be stopped early (Figure S2). In practice, we used ProbMinHash3a because it is the fastest among all variants due to computational optimization. In addition, the performance gain of ProbMinHash3a is achieved by calculating signature components (or sketch) not independently, but collectively^21^. To further speedup distance computation, we employed Simple Instruction Multiple Data (SIMD) for hamming distance computation between sketch vectors, which are collision probability of hashes. The SIMD distance implementation showed significant improvement only for large sketch sizes (e.g., > 1000).

### Link Prediction task

Link prediction predicts potential links between nodes in a network. In this task, we use the same setting as in Yang et.al. (2019)^9^. Specifically, we first randomly sample 20% of the edges from each graph as testing data, and then use the rest of the graph for training (Figure 1). After learning the node embeddings based on the training graph, we predict the missing edges by generating a ranked list of potential edges. For each pair of nodes, we use the Cosine or J_p_ (according to the embedding techniques) of their embeddings to generate the ranked list. We randomly sample 0.1% pairs of nodes for evaluation (0.01% on YouTube). We report the average precision@100 and recall@100 on the Blog, PPI, Wiki and DBLP datasets, and precision@1000 and recall@1000 on the YouTube dataset from 5 repeated trials. We did not benchmark GraphEmbed with large and sparse graph for link prediction tasks due to computational challenges (due to their large sizes).

### Multi-label Node classification tasks

Multi-label node classification tasks predict the most probable label(s) for some nodes based on other labeled nodes. In this experiment, we randomly pick a set of nodes as labeled nodes for training and use the rest for testing. To fairly compare node embeddings with different similarity measures, we train a support vector machine (SVM) classifier (cosine or J_p_ according to the embedding techniques) to return the most probable labels for each node. In classification tasks, we report the average Macro-F1 and Micro-F1 scores from 5 repeated trials, with 90% training ratio on the BlogCatalog, PPI, Wiki and DBLP datasets, and 10% training ratio on the YouTube dataset. In multi-label classification tasks, F1 score is defined below: 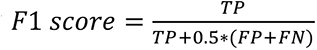.

We calculate the F1 score for each class in a One-vs-Rest (OvR) approach instead of a single overall F1 score, as seen in binary classification. The macro F1 score is computed using the arithmetic mean (aka unweighted mean) of all the per-class F1 scores while Micro F1 computes a global average F1 score by counting the sums of the True Positives (TP), False Negatives (FN), and False Positives (FP). We did not benchmark node classification tasks for large and sparse network datasets because training SVM classifiers for millions of nodes is computationally prohibitive. For weighted networks, the training and testing steps are the same but we only use the Coauth-MAG and Colisten-Spotify dataset. The calculation of Macro F1 and Micro F1 was performed via the Python package Scikit-Learn^70^.

### Other network embedding algorithms and parameters for benchmark

We chose a total of 15 published network embedding algorithms for benchmark. For random-walk based and matrix factorization-based algorithms, we choose DeepWalk^3^, LINE^6^, Node2Vec^4^, VERSE, NetMF^10^, GraRep^8^ and original HOPE^9^. Same parameters were used as in Yang et.al. (2019)^9^ for those algorithms. For hashing-based algorithms, we chose NetHash^19^, KatzSketch^9^ and original NodeSketch^18^. Same parameters were used for those algorithms. For randomized SVD-based network embedding algorithms, we chose RandNE^71^, FastRP^15^, ProNE^72^, LightNE^16^ and SkethNE^45^. The same parameters were used as in Xie et.al. (2023)^45^ for those algorithms.

### Parameter setup in GraphEmbed

For our HOPE implementation, a default setup was used according to original paper. However, for NodeSketch, decay weight ⍺ = 0.001 and number of iterations 5 for mullti-node classification tasks, and ⍺ = 0.3 and k = 5 for link prediction tasks were used. We varied ⍺ and k on several small to medium testing datasets to determine ⍺ and k according to the highest precision and recall (Figure S3A and B).

### Benchmark Datasets and Implementation Details

Several standard network benchmark datasets were used, including both undirected, directed, unweighted and weighted networks. Several large-scale networks were also included for testing the performance of GraphEmbed. See Table S1 for detailed characteristics of each dataset.

GraphEmbed utilizes compressed raw representation (CSR) format to represent adjacent matrix extracted from networks. The whole embedding system is in pure Rust and is fully parallelized whenever possible. Intel MKL library (OpenMP for multi-threading) or OpenBLAS (OpenMP for multi-threading) as the backend system level support for matrix operation through ndarray-linalg interface, the former is only supported on x86 systems while the latter supports various systems including ARM. All benchmarks throughout the paper were performed on 2 AMD EPYC 7302 16- core processor with a total of 64 threads available, unless specified otherwise. The output of embeddings is stored in MongoDB bson format (or binary JSON) for space efficiency and fast parsing. We provide scripts for parsing embeddings so that the embeddings can be easily used in Python Numpy vector or matrix format.

## Code and Data Availability

Code for GraphEmbed can be found here: https://gitlab.com/Jianshu_Zhao/graphembed or via Zenodo^73^. ProbMinHash library can be found here: https://github.com/jean-pierreBoth/probminhash. Scripts for reproducing results can be found here: https://gitlab.com/Jianshu_Zhao/graphembed-analysis. All testing data used in this study are open sourced and can be found freely online. GraphEmbed is also available in Bioconda (https://anaconda.org/bioconda/graphembed) or PyPi (https://pypi.org/project/graphembed-rs/) as a python library, also as part of the scikit-bio library.

## Acknowledgements

We want to thank Daniel McDonald and Igor Sfiligoi for the helpful discussion about the main results of our manuscript. We want to thank Ertl Otmar for reference implementation of ProbMinHash algorithms. We also want to thank Claire Wang and Sherlyn Weng for testing GraphEmbed, and Celeste Allaband for help with proofreading. This work was funded in part by the Department of Energy, USA to R.K. (DE-SC0024320)

## Author contribution

J.Z, J.P.B and R.K designed the work. J.Z and J.P.B implemented the ProbMinHash and GraphEmbed library. R.K supervised the work. J.Z. did the analysis and benchmark. J.Z., J.P.B and R.K wrote the manuscript. All authors read and edited the manuscript.

